# Nanopore Adaptive Sampling Enriches for Antimicrobial Resistance Genes in Microbial Communities

**DOI:** 10.1101/2023.06.27.546783

**Authors:** Danielle C. Wrenn, Devin M. Drown

**Affiliations:** Department of Biology and Wildlife, University of Alaska Fairbanks, Fairbanks, Alaska, USA; Institute of Arctic Biology, University of Alaska Fairbanks, Fairbanks, Alaska, USA

**Keywords:** antibiotic resistance, environmental microbial communities

## Abstract

Antimicrobial resistance (AMR) is a global public health threat. Environmental microbial communities act as reservoirs for AMR, containing genes associated with resistance, their precursors, and the selective pressures to encourage their persistence. Genomic surveillance could provide insight into how these reservoirs are changing and their impact on public health. The ability to enrich for AMR genomic signatures in complex microbial communities would strengthen surveillance efforts and reduce time-to-answer. Here, we test the ability of nanopore sequencing and adaptive sampling to enrich for AMR genes in a mock community of environmental origin. Our setup implemented the MinION mk1B, an NVIDIA Jetson Xavier GPU, and flongle flow cells. We observed consistent enrichment by composition when using adaptive sampling. On average, adaptive sampling resulted in a target composition that was 4x higher than a treatment without adaptive sampling. Despite a decrease in total sequencing output, the use of adaptive sampling increased target yield in most replicates.

## Introduction

Antimicrobial resistance (AMR) is a public health threat of great magnitude, accounting for over 2.8 million infections and 35,000 deaths annually in the U.S. alone [1]. Resistant pathogens pose the most direct risk to human health. However, the AMR genes present in pathogens only represent a small proportion of a much larger collection of AMR genes, the antibiotic resistome. As described by Wright [2] and D’Costa et al. [3], the antibiotic resistome includes all antimicrobial resistance genes and their precursors, the majority of which reside in nonpathogenic microbial communities.

Environmental microbial communities serve as important contributors to the resistome. They are dynamic reservoirs where a variety of factors influence the evolution, exchange, and persistence of genes that confer resistance. Resistance mechanisms originated in the environment [2][4] where production of and resistance to antimicrobial agents assists microorganisms in the battle for territory and resources [5][4]. External influences like human and agricultural waste streams introduce resistant organisms, resistance genes, and pharmaceutical antimicrobial agents into the environment [6][7]. Once within the community these agents provide additional selective pressure for resistance and the genes provide new material for exchange.

The exchange of resistance genes between community members continues even in the absence of selective pressure [8]. This continued exchange is likely one reason why antimicrobial resistance can persist even after the removal or reduction of antimicrobial exposure [9][10]. The likelihood of the reversal of a resistant community to a susceptible one is a complex landscape influenced by mutation rate, fitness cost, and compensatory evolution [11].

AMR genes can then be shared from environmental microbial communities to human and animal pathogens [12][13]. A One Health approach, one that recognizes the interconnection of human, animal, and environmental health, has grown in popularity regarding microbiology and AMR research [14][15][16][17]. The World Health Organization [18] and the CDC [1] have both endorsed a One Health approach as an effective strategy for addressing AMR.

Despite the increased interest in investigating environmental AMR, gaps in knowledge still exist regarding the exchange of AMR genes between environmental organisms and pathogenic communities, the effects of abiotic factors on the persistence and evolution of environmental AMR, and the effects of clinical and agricultural interventions on environmental microbial communities. Genomic surveillance of genes associated with AMR could provide important insight into how these dynamic reservoirs are impacting public health. Genomic surveillance allows for the monitoring of the entire resistome, that is AMR genes both inside and outside of pathogenic organisms as well as their precursors. It also allows for the detection of “silent” AMR genes, those present in susceptible organisms but that may confer resistance following a shift in host or environment. However, genomic sequencing is typically time, resource, and cost-intensive, especially outside of clinical settings.

Nanopore sequencing presents an opportunity for the development of a cost-effective and portable genomic surveillance tool. While more commonly used sequencing technologies sequence via DNA synthesis, nanopore sequencing determines genetic sequences by detecting a change in current as DNA strands are pulled through nanopores on the flow cell [19]. The technology allows for a streamlined, resource conservative library preparation. It also allows for unique features like adaptive sampling [20][21]. Traditional sequencing technologies, such as Illumina, achieve enrichment by using reactions such as PCR prior to sequencing. Pre-sequencing enrichment necessitates the use of additional time and resources including synthesized primers. In contrast, adaptive sampling requires no change in library preparation as it leverages the ability of each nanopore to independently accept and reject strands of DNA during sequencing. The enrichment or depletion of user-defined targets is therefore achieved entirely *in silico*, without the need for additional time, resources, or effort. The MinION, the smallest genomic sequencer currently commercially available, boasts incredible portability (with minimal power consumption) in addition to being capable of adaptive sampling.

Other studies have utilized nanopore sequencing and the adaptive sampling feature for the detection of AMR genes in clinical samples through both host depletion [22] and AMR gene enrichment [23]. However, the exploration of adaptive sampling for the enrichment of AMR genes in environmental metagenomic samples is limited. Our goal is the development of a novel toolbox that is optimized for the rapid, resource conservative surveillance of AMR associated genes in environmental microbial communities. The principal question of this study is whether adaptive sampling is capable of enriching (by composition) for AMR associated genes in a mock community of environmental origin. This study investigated metrics of performance including enrichment by target yield and the proportion of panel that was successfully detected.

## Methods

### Experimental Design

To test the effects of adaptive sampling on AMR gene enrichment, we included two treatments: adaptive sampling on and off. We simultaneously implemented these two treatments by turning on adaptive sampling for 50% of the sequencing nanopores on the flow cell. The other half of the flow cell sequenced the library using the traditional, non-selective, method (adaptive sampling off). With this design, we were able to control for variability in our library preparation from run to run.

We generated a mock community from bacterial isolates with known AMR genes from previously isolated and archived soil samples from the Fairbanks Permafrost Experiment Station [24]. Original bacterial culturing and isolation methods are described in Haan and Drown [24]. We selected six community members (TH25, TH28, TH41, TH57, TH79, TH81) representing five genera (*Serratia, Bacillus, Erwinia, Pantoea*, and *Pseudomonas*) of common soil bacteria associated with permafrost thaw to compose the final mock community. These members were selected to include phylogenetic diversity that would include a diverse set of AMR genes. For this experiment, we extracted and purified DNA from previously frozen cells using the DNeasy UltraClean Microbial Kit (Qiagen) according to manufacturer instructions. After quantification of DNA concentration from extractions using a Qubit (Thermo Fisher Scientific), we pooled all members of the community by equal mass (1000 ng).

Using the published sequences (Biosample accessions SAMN17054805, SAMN17054834, SAMN17054856, SAMN09840060, SAMN17054818, SAMN17054803) [24], we identified all AMR gene regions using the Resistance Gene Identifier (RGI) version 5.1.0 and the Comprehensive Antibiotic Resistance Database (CARD) [25] version 3.0.9 to compose the target gene panel using exclusively strict and perfect hits. Targeted genes and the number of gene copies per community member are specified in Appendix A. The expansion of targeted regions through the inclusion of flanking DNA has been implemented by previous studies [26][20] and is recommended by Oxford Nanopore to increase target output. We used a custom script to expand the gene region and include a flanking region of DNA in each targeted region. Each flanking region was the size of the prepared library’s N50 (5075 bp). We extracted target sequences using Geneious Prime 2022.1.1 (https://www.geneious.com). The resulting multi-fasta file contained 52 unique sequences and served as the adaptive sampling reference.

### Library Preparation & Sequencing

We used the Oxford Nanopore Technologies (ONT) Rapid Sequencing Kit (SQK-RAD004) for sequencing library preparation. For each library, we used 200 ng of input DNA from our mock community. We followed the manufacturer protocol except for the exclusion of the bead cleanup and Qubit quantification steps to maximize the DNA quantity carried forward into sequencing.

The MinION mk1B, an NVIDIA Jetson Xavier GPU, and flongle flow cells (FLO-FLG001, R9.4.1) were used for sequencing. We configured the Xavier GPU with MinKNOW (MinKNOW Core version 4.5.4) following instructions from Benton (2022) [27]. All sequencing runs lasted eight hours. Half of the flow cell (63 channels) was designated for adaptive sampling; the other half sequenced normally (adaptive sampling off). We alternated the side of the flow cell performing each treatment for each replicate. Each flow cell was used twice (technical replicates). We completed thirteen total sequencing runs.

### Data & Statistical Analysis & Visualization

We used Guppy version 6.1.3 to base call the raw sequencing data using the super-accuracy model (dna_r9.4.1_450bps_sup.cfg) and filtered by minimum quality score (Q score ≥ 10). During adaptive sampling, the first 500-1000 bp of a template strand of DNA is sequenced. Regardless of the adaptive sampling algorithm decision (accept or reject), that preliminary sequence is output. To remove these very short reads, we filtered the output by length (>1000 bp) using seqtk version 1.3 [28]. We aligned the filtered output to the community metagenome with Minimap2 version 2.22 [29] using the Oxford Nanopore genomic reads preset (-ax map-ont). We used SAMtools version 1.15.1 to exclude supplementary and secondary alignments (-F 2308) [30]. We used the sequencing summary generated by Guppy to calculate the average number of active pores. We first subset the data by treatment, then binned the data into one-hour intervals. The number of unique channels generating reads were then calculated for each hour and then averaged across the length of the run.

We calculated the summary statistics for each run (yield, mean quality score) using NanoStat version 1.6.0 [31]. To calculate target yield, we used SAMtools coverage and depth. SAMtools coverage was implemented to determine the number of reads that contained targeted AMR regions. Depth was used to determine the number of nucleotides that aligned to targeted AMR regions. In these calculations, AMR regions refer to the AMR genes without expanded flanking regions. To avoid a single read being counted multiple times in our target yield calculations, we only included unique alignments in downstream analysis.

We utilized base R (version 4.2.2) and the R car package [32] for statistical analysis. We used the Shapiro-Wilk test to determine data normality. Variance homogeneity was determined using either an F-test or Levene’s test as appropriate. A Two Sample t-test, Welch’s T-test, or Wilcoxon signed-rank test was then employed to evaluate the significance of any difference between treatments (α = 0.05). For data visualization, we used ggplot2 [33].

## Results

Over the course of 4 days, we completed 13 sequencing runs. We excluded three runs that lacked available pores at the end of the first technical replicate resulting in 10 sequencing runs, including technical replicates, used for analysis (Table 1). Our maximum output run generated over 281 Mb of data, the lowest over 19 Mb. On average, sequencing runs yielded 103,728,356 bp and contained 42,176,321 bp after filtering by quality and length. On average, second technical replicates generated 62% less data (*μ*_first_ = 150.6 Mbases, *μ*_second_ = 56.86 Mbases), and lower mean output quality (decrease of 12.7%). Filtering by quality and length resulted in a 59% decrease in yield but a 23% increase in quality (Table 1). Only post-filtering data were used in alignment and target yield quantification.

**Table 1.** Sequencing output metrics prior to and following filtering for quality and read length.

|  |  | Pre-Filtering |  | Post-Filtering |  |
| --- | --- | --- | --- | --- | --- |
| Run | Technical Replicate | Yield (bp) | Mean Quality (Q) Score | Yield (bp) | Mean Quality (Q) Score |
| 1 | 1 | 281,215,264 | 10.3 | 109,639,119 | 12.3 |
| 2 | 2 | 98,850,455 | 8.2 | 18,970,458 | 11.4 |
| 3 | 1 | 170,831,418 | 10.6 | 86,179,378 | 12.5 |
| 4 | 2 | 77,078,419 | 9.4 | 22,621,178 | 12.2 |
| 5 | 1 | 99,711,784 | 10.5 | 40,575,094 | 12.8 |
| 6 | 2 | 51,020,783 | 9.1 | 20,600,281 | 12.5 |
| 7 | 1 | 146,519,076 | 11.3 | 62,208,499 | 13.3 |
| 8 | 2 | 38,318,184 | 10.8 | 21,961,623 | 12.5 |
| 9 | 1 | 54,700,216 | 11.5 | 32,243,307 | 12.8 |
| 10 | 2 | 19,037,961 | 9.8 | 6,764,271 | 12.0 |

Regardless of sequence identity, we observed a significant decrease in sequencing output when using adaptive sampling (*t* = -6.67, *p* = 2.968e^-6^) (Figure 1). Although the adaptive sampling off treatment appeared to display greater variability in output between runs (σ^2^ = 1.09) compared to when adaptive sampling was turned on (σ^2^ = 0.42), this difference was not statically significant (F = 0.385, *p* = 0.171). Here, sequencing output refers to the total sequencing yield (pre-filtering) per treatment. While we split the flow cell evenly across treatments, there may still exist some variation in pore availability between flow cells and treatments. To control for this variation, we normalized these yields by the average number of active pores during the sequencing run. The need for this normalization was compounded by our use of technical replicates where we saw an increase in variation of active pores.

**Figure 1.**
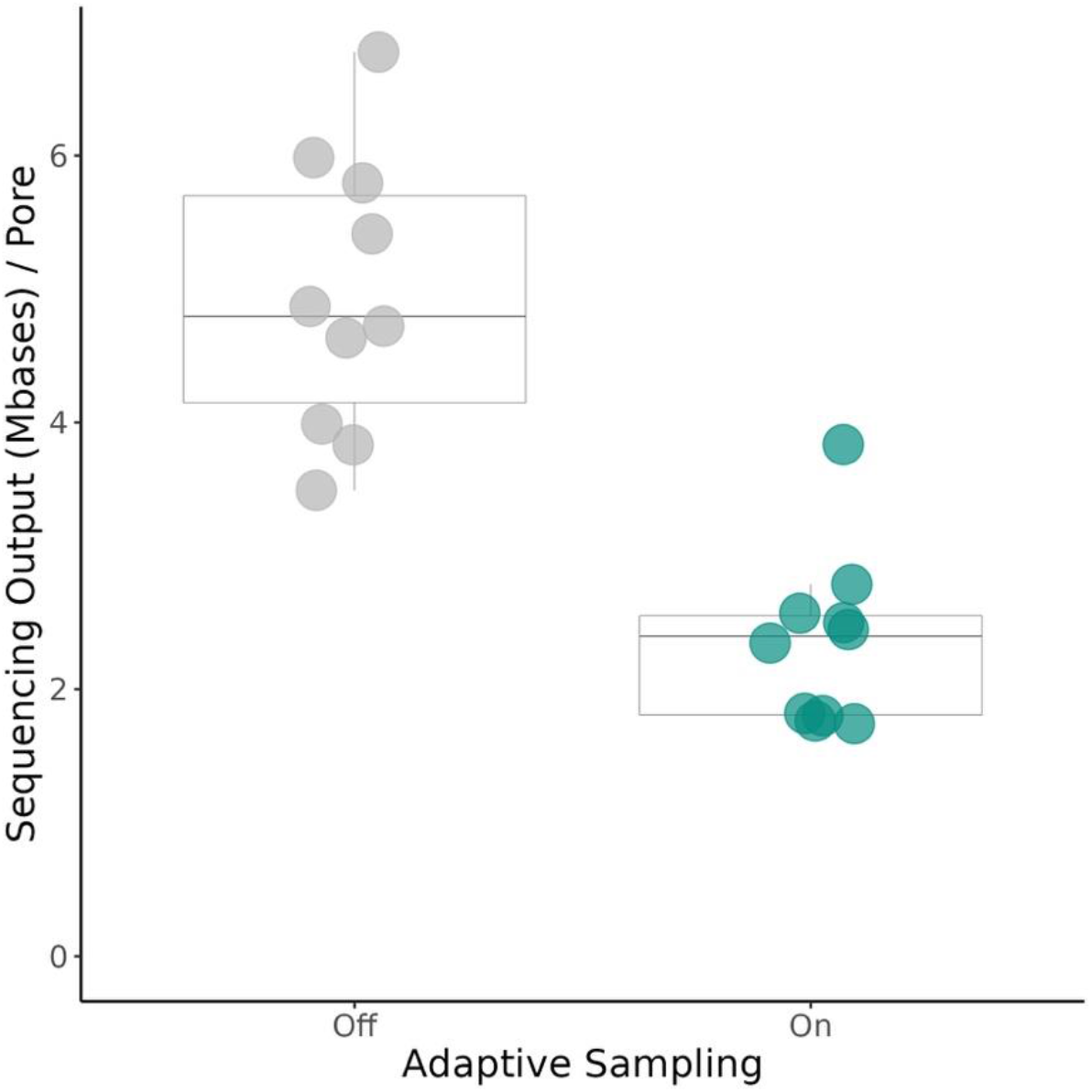
A comparison of total sequencing output with and without the use of adaptive sampling. Sequencing output refers to all the data generated prior to filtering for quality and length. Total output is normalized using the average number of active pores during the entire run duration for each treatment. Statistical analysis used a paired Welch’s T-test (*t* = -6.67, *p* = 2.968e^-6^). *μ*_OFF_ = 4.95 Mb, *μ*_ON_ = 2.36 Mb (n = 10)

Next, we evaluated AMR gene target enrichment by composition. This is a measure of the fraction of the sequencing output that includes the targeted AMR genes. To do this, we calculated the percent target composition for each treatment and sequencing run where percent composition was calculated as follows: (Output aligned to target AMR genes (bp) / Total pre-filtering sequencing output (bp)) * 100. Despite decreased yield observed in the adaptive sampling treatment (Figure 1), the proportion of sequencing output that is composed of target AMR genes is significantly greater for the adaptive sampling treatment (V = 55, *p* = 0.002). On average, percent target composition achieved by adaptive sampling was over 4x higher than that observed in the control treatment (Figure 2). We found that over 0.42% of the output of the adaptive sampling treatment represented the target gene sequences, on average. For context, we estimate that the true representation of the targeted AMR genes in our sample metagenome is 0.24%.

**Figure 2.**
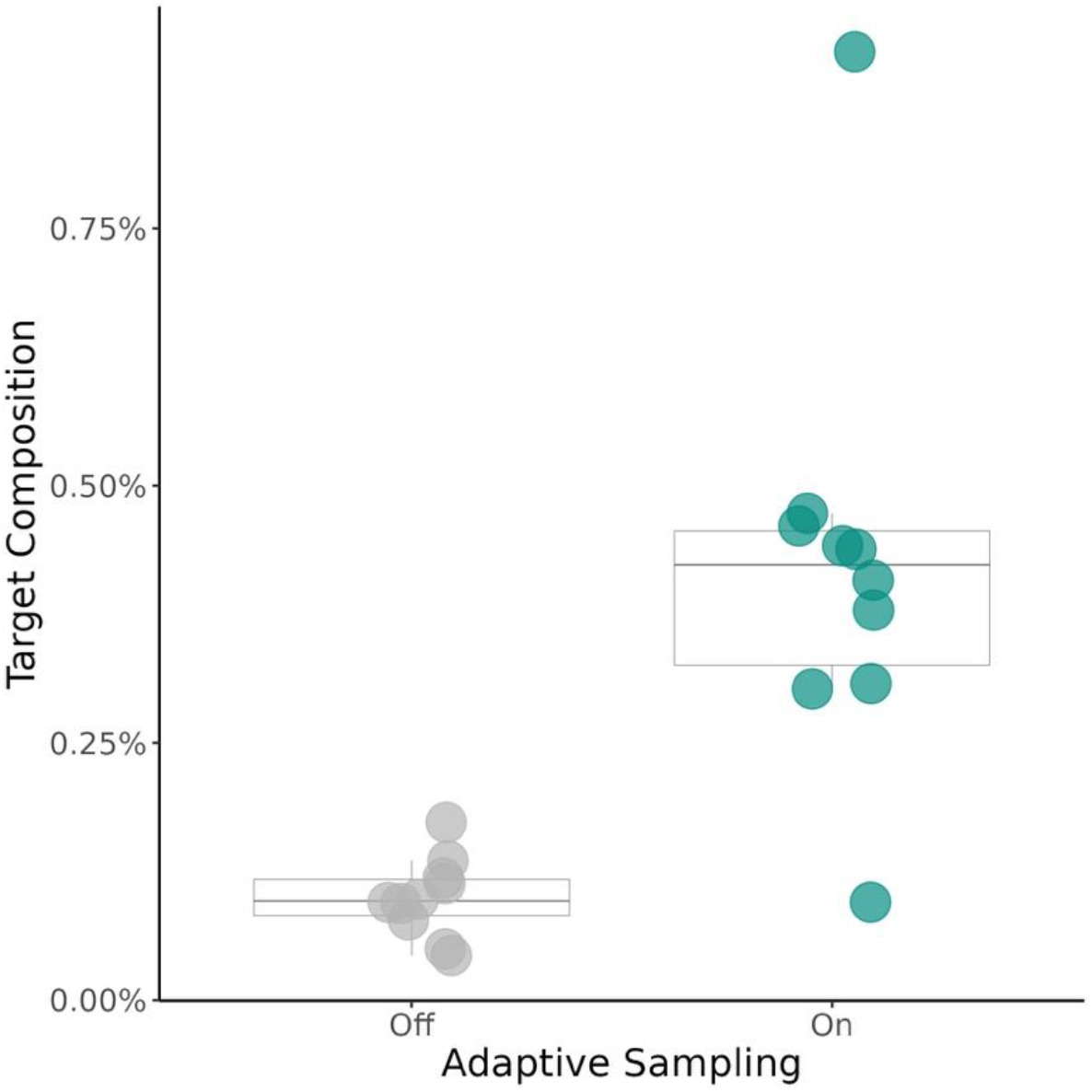
Comparison of the target composition of total sequencing output with and without the use of adaptive sampling. Percent target composition was calculated as the output aligned to a targeted AMR region (bp) / total sequencing output (pre-filtering) (bp) * 100. Statistical analysis used a Wilcoxon signed-rank test (V = 55, *p* = 0.002). *μ*_OFF_ = 0.1%, *μ*_ON_ = 0.42% (n = 10)

We also evaluated enrichment by target yield. This is a measure of the sequencing yield (Kbases) that was solely composed of the designated target genes. To measure the performance difference between treatments, we calculated the percent difference between treatments for each of our sequencing runs as normalized by the control run. Percent difference was calculated as follows: [(target yield (bp) per average active pores with adaptive sampling - target yield (bp) per average active pores without adaptive sampling) / target yield (bp) per average active pores without adaptive sampling] * 100. Values above zero indicated that the use of adaptive sampling resulted in greater target yield. The difference in target yield is significantly greater than zero (V = 54, *p* = 0.00195) (Figure 3). Adaptive sampling outperformed the control treatment in this metric for nine out of ten replicates. The mean percent difference between the two treatments was 104.6% representing a greater than two-fold increase in target yield when adaptive sampling was used (Figure 3).

**Figure 3.**
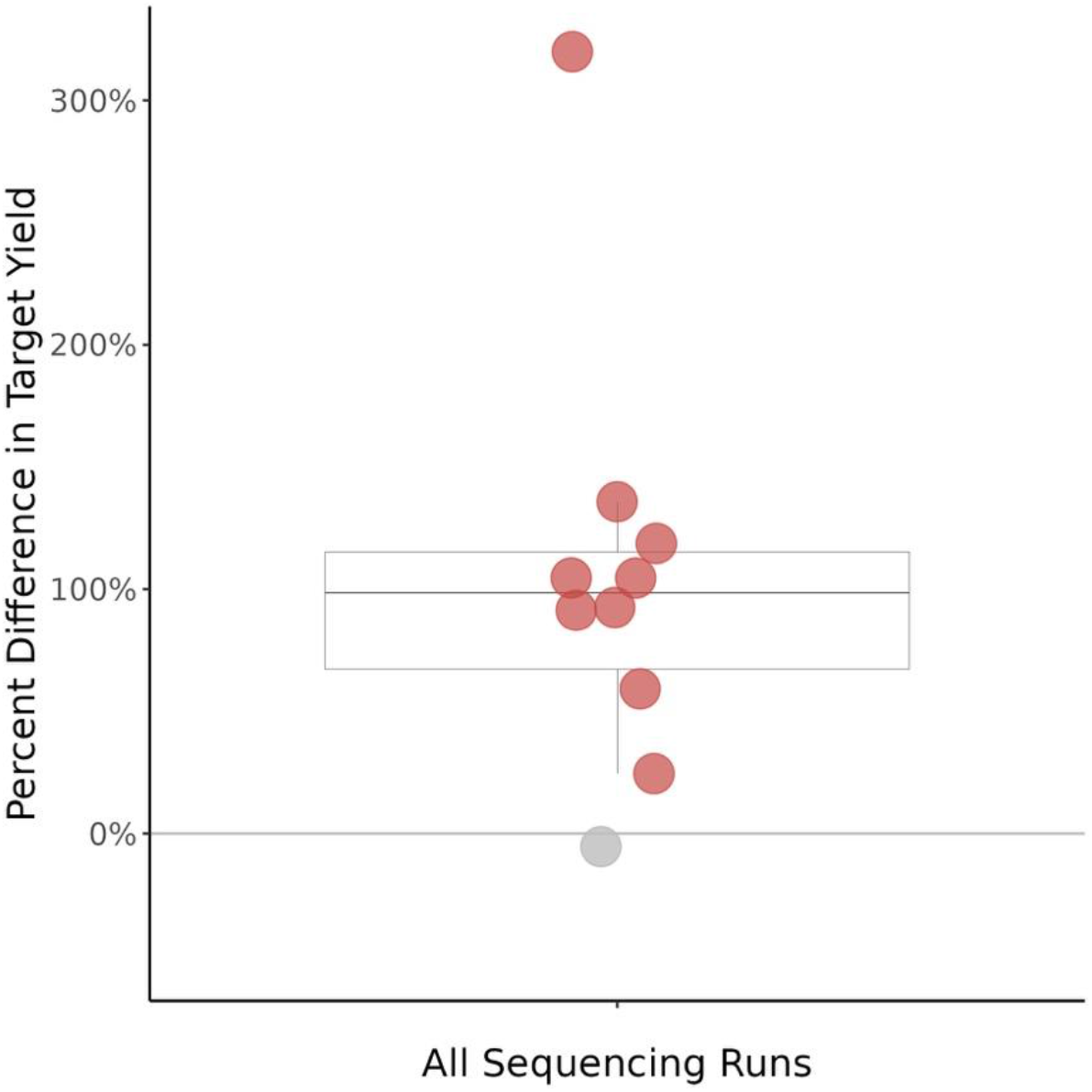
The percent difference in target yield between adaptive sampling on and adaptive sampling off sides of each flow cell. Percent difference was calculated with the half of the flow cell sequencing normally (adaptive sampling off) as the initial value. Red points denote a difference > 0%, gray points denote a difference ≤ 0%. Statistical analysis used a Wilcoxon signed-rank test (V = 54, *p* = 0.00195). *μ* = 104.6% (n = 10)

Finally, we looked at the proportion of our target panel that was detected by each treatment. Our criteria for detection were as follows: 100% coverage of the AMR region with a minimum depth of 2 nucleotides at every position. Due to output requirements inherent in the criteria, sequencing runs that generated less than 25 Mb of post-filtering data were excluded in this analysis (n = 5). When adaptive sampling was used 21.9% of the panel was detected, on average. This is more than double the average 8% observed when adaptive sampling was not in use (Figure 4). The maximum proportion detected was 36.5% and 21.2% with adaptive sampling on and off respectively. Within a sequencing run, the side of the flow cell implementing adaptive sampling consistently detected more of the panel than its non-adaptive sampling counterpart; however, we did not find the difference between these two treatments to be significant (V = 15, *p* = 0.0625).

**Figure 4.**
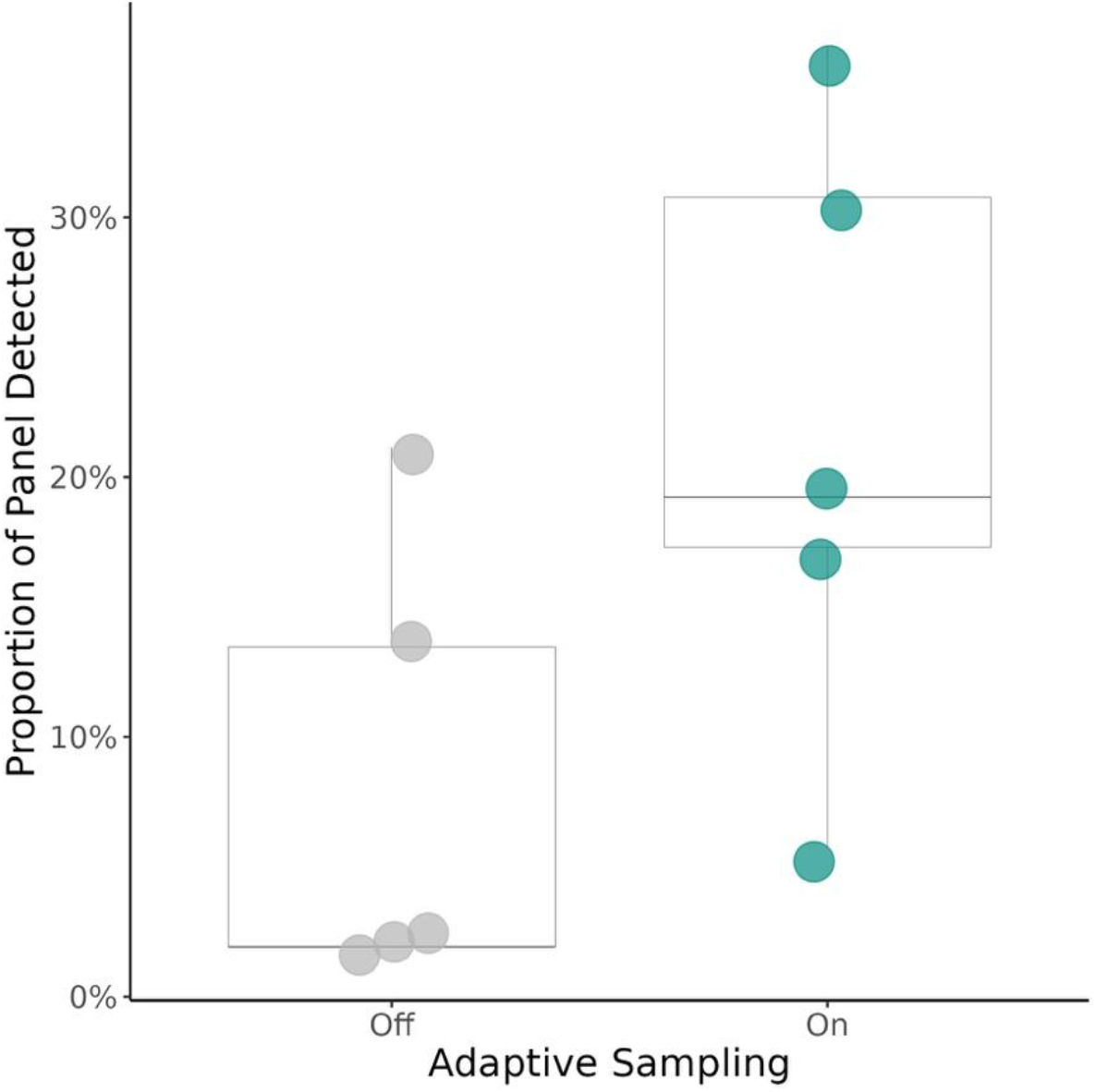
Proportion of the target AMR gene panel detected for adaptive sampling on and off treatments. A successful detection was defined as 100% AMR gene coverage with ≥ 2bp depth at every position. Statistical analysis used the Wilcoxon signed-rank test (V = 15, *p* = 0.0625). *μ*_OFF_ = 8.1%, *μ*_ON_ = 21.9% (n = 5)

## Discussion

This research represents the first steps in the development of a novel toolbox for the rapid, resource conservative surveillance of AMR associated genes in environmental microbial communities. Our goal in this study was the evaluation of adaptive sampling’s ability to enrich (by composition) for AMR associated genes in a known sample. We found that adaptive sampling can enrich for antimicrobial resistance genes in our mock community. We observed consistent enrichment by composition when using adaptive sampling regardless of overall sequencing yield.

Martin et al. [34] demonstrated the ability of adaptive sampling to enrich by composition for genomes in metagenomic samples, here we have demonstrated that adaptive sampling can enrich for much smaller targets - i.e., AMR genes in microbial communities. Our observations regarding enrichment by target yield are encouraging. Other studies have noted the association between enrichment by yield and sequencing run output [34]. This is due, in part, to the variability in pore quality and pore loss between flow cells. Our use of technical replicates, where second technical replicates began with fewer available pores and those that remained were likely decreased in quality, may have further exacerbated this effect in our study.

Further optimization could increase enrichment by yield using adaptive sampling. The available literature suggests that template length, target size, percent identity, and the above-mentioned pore availability can all impact enrichment by yield [23][34]. The ratio of target size to template length contributes to the likelihood that the pore sees that the target sequence is present before the algorithm rejects that strand. Small targets on long templates have a higher likelihood of being missed. The lower the percent identity between the target and template, also increases the likelihood that a sequence will not be recognized as on-target [23]. This is due to adaptive sampling’s reliance on live alignment of template strands to target sequence data to determine target presence. Finally, pore quality and availability directly impact the sequencer’s ability to generate both on-target and off-target data [34].

For these experiments, we relied on low cost flongle flow cells that cost a fraction of the cost of a traditional flow cell while generating a fraction of the yield. However, low target yield and low sequencing output overall, contributed to the inability of either treatment to detect greater than 37% of our target panel. Optimization that improves target yield may also improve panel detection. We employed the use of increased target size in the pursuit of greater enrichment by yield. Further work is warranted to explore the employment of other strategies to produce consistent enrichment by target yield in our protocol.

The expansion of current knowledge regarding resistance in environmental microbial communities benefits a One Health approach to addressing the threat of AMR. Environmental microbial communities play an important role in the origin, persistence, and dissemination of resistance mechanisms [2][4][9][10]. The MinION, with its incredible portability and ability to perform adaptive sampling, could reduce time-to-answer and economic barriers to genomic surveillance of environmental reservoirs of AMR associated genes.

Other studies have described the potential benefit of using adaptive sampling for the reduction of time-to-diagnosis in clinical samples [22]. Reduced time-to-answer in an environmental context could allow for better informed preventative public health action, industry standard modification, and policy implementation. The scope of this study was limited in terms of communities tested and AMR genes targeted. However, its results are promising for the development of a flexible, portable, and cost-effective AMR surveillance tool. Future work could include the expansion of the target gene panel so that the toolbox could be applied to a larger cohort of microbial communities and the testing of the protocol on an environmental sample.

## Availability of Test Data

The data set supporting the results of this article is available in the SRA database under BioProject PRJNA982864.

## List of Abbreviations

AMR: antimicrobial resistance
bp: base pair
CDC: Center for Disease Control and Prevention
Kbase: kilobase
Mbase: megabase
ONT: Oxford Nanopore Technologies
WHO: World Health Organization

## Declarations

## Ethical Approval

Not Applicable.

## Author’s Contributions

Conceptualization, Investigation, Formal Analysis, Software, Methodology, Validation, Data Curation, Resources, Funding Acquisition, Visualization: DCW, DMD; Writing - Original Draft Preparation: DCW; Writing - Review & Editing: DCW, DMD.

## Acknowledgements

We would like to thank Tracie Haan for supplying the isolates used in our mock community and technical support. We would like to thank Upasana Arora, Jeremy Buttler, Bevyn Cover, Ursel Schütte, and Jorda Kovash for their constructive feedback. We would like to thank Miles Benton who kindly provided detailed instructions for computer setup and inspiration for this project. We acknowledge the generous support of the Institute of Arctic Biology and Logan Mullen in the IAB Genomic Core Laboratory.

## Appendix A. Targeted Genes

| Member |  | TH25 | TH28 | TH41 | TH57 | TH79 | TH81 |
| --- | --- | --- | --- | --- | --- | --- | --- |
| Genus |  | *Bacillus* | *Serratia* | *Pseudomonas* | *Bacillus* | *Erwinia* | *Pantoea* |
|  | AMR Gene |  |  |  |  |  |  |
|  | BcII | 1 | 0 | 0 | 1 | 0 | 0 |
|  | FosB | 1 | 0 | 0 | 1 | 0 | 0 |
|  | MCR-4.5 | 1 | 0 | 0 | 0 | 0 | 0 |
|  | tet(45) | 1 | 0 | 0 | 0 | 0 | 0 |
|  | CRP | 0 | 1 | 0 | 0 | 1 | 1 |
|  | Escherichia coli EF-Tu mutants conferring resistance to Pulvomycin | 0 | 1 | 0 | 0 | 0 | 0 |
|  | Haemophilus influenzae PBP3 conferring resistance to beta-lactam antibiotics | 0 | 1 | 0 | 0 | 1 | 1 |
|  | Klebsiella pneumoniae KpnF | 0 | 1 | 0 | 0 | 1 | 1 |
|  | Klebsiella pneumoniae KpnH | 0 | 1 | 0 | 0 | 1 | 1 |
|  | adeF | 0 | 5 | 3 | 0 | 2 | 2 |
|  | emrR | 0 | 1 | 0 | 0 | 1 | 1 |
|  | msbA | 0 | 1 | 0 | 0 | 1 | 1 |
|  | Acinetobacter baumannii AbaQ | 0 | 0 | 1 | 0 | 0 | 0 |
|  | Pseudomonas aeruginosa soxR | 0 | 0 | 1 | 0 | 0 | 0 |
|  | armA | 0 | 0 | 1 | 0 | 0 | 0 |
|  | MCR-4.1 | 0 | 0 | 0 | 1 | 0 | 0 |
|  | sgm | 0 | 0 | 0 | 1 | 0 | 0 |
|  | CARB-23 | 0 | 0 | 0 | 0 | 1 | 0 |
|  | Escherichia coli ampH beta-lactamase | 0 | 0 | 0 | 0 | 1 | 1 |
|  | Morganella morganii gyrB conferring resistance to fluoroquinolone | 0 | 0 | 0 | 0 | 1 | 1 |
|  | PmrF | 0 | 0 | 0 | 0 | 1 | 0 |
|  | BES-1 | 0 | 0 | 0 | 0 | 0 | 1 |
|  | Escherichia coli UhpT with mutation conferring resistance to fosfomycin | 0 | 0 | 0 | 0 | 0 | 1 |
|  | Klebsiella pneumoniae KpnE | 0 | 0 | 0 | 0 | 0 | 1 |
|  | amrB | 0 | 0 | 0 | 0 | 0 | 1 |

